# Neurofibromin regulates metabolic rate via neuronal mechanisms in *Drosophila*

**DOI:** 10.1101/834788

**Authors:** Valentina Botero, Bethany A. Stahl, Eliza C. Grenci, Tamara Boto, Scarlet J. Park, Lanikea B. King, Keith R. Murphy, Kenneth J. Colodner, James A. Walker, Alex C. Keene, William W. Ja, Seth M. Tomchik

## Abstract

Neurofibromatosis type 1 (NF1) is a genetic disorder predisposing patients to a range of features, the most characteristic of which include areas of abnormal skin pigmentation and benign tumors associated with peripheral nerves, termed neurofibromas. Less common, but more serious symptoms also include malignant peripheral nerve sheath tumors, other malignancies, and learning disabilities. The *NF1* gene encodes neurofibromin, a large protein that functions as a negative regulator of Ras signaling and mediates pleiotropic cellular and organismal function. Recent evidence suggests NF1 may regulate metabolism, though the mechanisms are unknown. Here we show that the *Drosophila* ortholog of NF1, dNf1 regulates metabolic homeostasis in fruit flies by functioning within a discrete brain circuit. Loss of dNf1 increases metabolic rate and feeding, enhances starvation susceptibility, and decreases lipid stores while increasing lipid turnover rate. The increase in metabolic rate is independent of locomotor activity (grooming), and maps to a subset of neurons in the ventral nervous system. The feeding and metabolic rate effects are due to loss of dNf1 in the same set of neurons, suggesting that increased feeding may be a compensatory effect driven by the increase in metabolic rate and lipid turnover. Finally, we show that the Ras GAP-related domain of neurofibromin is required for normal metabolism, demonstrating that Ras signaling downstream of dNf1 mediates the metabolic effects. These data demonstrate that dNf1 regulates metabolic rate via neuronal mechanisms, suggest that cellular and systemic metabolic alterations may represent a pathophysiological mechanism in NF1, and provide a platform for investigating the cellular role of neurofibromin in metabolic homeostasis.

## INTRODUCTION

Neurofibromatosis type 1 (NF1) is a genetic disorder affecting 1 in ~3,500 individuals. Caused by mutations in the *NF1* gene, this disorder is characterized by benign tumors of the nervous system called neurofibromas, as well as an increased susceptibility to a range of complications including various cancers and neurocognitive deficits (e.g., attention-deficit/hyperactivity disorder, autism spectrum disorder, visuospatial memory impairments) (Diggs-Andrews and Gutmann, 2013; Hyman et al., 2005). The *NF1* tumor suppressor gene encodes a large protein called neurofibromin, which contains a central GAP-related domain (GRD) that enhances the GTPase activity of the small guanine nucleotide binding protein Ras, thereby down-regulating its biological activity (Cichowski and Jacks, 2001; Ratner and Miller, 2015). Ras, in turn, signals through multiple effectors, including mTOR, ERK, and, potentially cAMP/PKA indirectly (Gehart et al., 2010). Due to the numerous cellular functions of Ras, as well as potential interactions of neurofibromin with other signaling molecules (Ho et al., 2007; Ratner and Miller, 2015; Xie et al., 2016), loss of NF1 results in pleiotropic effects on cellular and organismal physiology which underlie this multisystem disorder. A major question is what are the central pathophysiological mechanism(s) of NF1; how does loss of NF1 alter cellular physiology resulting in cellular/behavioral phenotypes seen in NF1 patients and in animal models for the disease?

Emerging evidence suggests that NF1 regulates cellular and organismal metabolism. NF1 has been associated with short stature, as well as reduced body mass index in males (Koga et al., 2016; Souza et al., 2016), although the mechanisms underlying these manifestations are unknown. Alterations in certain metabolites have been reported in NF1 patients, though these are sex-specific (Koga et al., 2016). The incidence of diabetes mellitus, deaths from diabetes mellitus, and fasting blood glucose levels are lower in individuals with NF1 than in healthy controls (Martins et al., 2016). Of potential importance for NF1-associated tumorigenesis, loss of NF1 increases glycolysis and decreases respiration via Ras/ERK signaling in mitochondria (Masgras et al., 2017). Finally, increased resting energy expenditure has been reported in females with NF1 (Souza et al., 2019). Therefore, NF1 has the potential to regulate cellular and organismal metabolism, and alterations in metabolism may contribute to its pathophysiology. However, clinical studies have produced sometimes contradictory results across cohorts/sexes, often include low sample sizes, and the multisystemic nature of the disorder precludes identification of the mechanism(s) underlying the reported effects. Thus, the fundamental 5effects of NF1 on metabolism remain an open question.

We have investigated the role of the *Drosophila melanogaster* ortholog of NF1 (dNf1) in the regulation of metabolism. *Drosophila* neurofibromin is ~60% identical to the human protein and similarly mediates Ras signaling (The et al., 1997; Walker et al., 2013). Flies with *dNf1* mutations exhibit small body size (The et al., 1997; Walker et al., 2013; Walker et al., 2006), impaired circadian rhythms (Williams et al., 2001), learning and memory deficits (Buchanan and Davis, 2010; Gouzi et al., 2011; Guo et al., 2000), decreased lifespan via increased susceptibility to oxidative stress (Tong et al., 2007), and increased spontaneous grooming (King et al., 2016). These changes demonstrate widespread alterations in cellular/neuronal function, raising the possibility that metabolic alterations could be a central phenotype of NF1 deficiency across phyla.

## RESULTS

### Loss of dNf1 increases metabolic rate via neuronal mechanisms

To test whether dNf1 affects metabolic rate, we examined CO_2_ production via respirometry (Fig. 1A) (Yatsenko et al., 2014). In insects, whole body CO_2_ production provides a readout of metabolic rate and can be readily measured in freely moving animals (Lighton, 2008). We first compared *Nf1*^*P1*^ mutants, which harbor a large deletion in the *dNf1* locus, including the central catalytic GRD (The et al., 1997), with *w*^CS10^ controls. *Nf1*^*P1*^ mutant adult flies exhibited significantly elevated CO_2_ output compared to controls (Fig. 1B and S1A), suggesting that dNf1 plays a role in metabolic regulation. For additional confirmation, we tested a heteroallelic combination of the *Nf1*^*P1*^ deletion with a nonsense mutation, *Nf1^E1^* (Walker et al., 2006). *Nf1^P1/E1^* mutants also exhibited significantly elevated CO_2_ output relative to *w*^CS10^ control flies (Fig. 1C and S1B). To test whether the effect observed in *dNf1* mutants is localized to neurons, we knocked down dNf1 with RNAi using the Gal4/UAS system. In independent experiments, two different pan-neuronal Gal4 drivers were used to drive the RNAi: nSyb-Gal4 and R57C10-Gal4. In each case, a significant elevation in CO_2_ production was observed in the experimental group relative to flies harboring either the heterozygous driver (Gal4/+) or effector (UAS/+) transgenes alone (Fig. 1D and S1C). This suggests that the loss of dNf1 function elevates metabolic rate via neuronal mechanisms.

**Figure 1.**
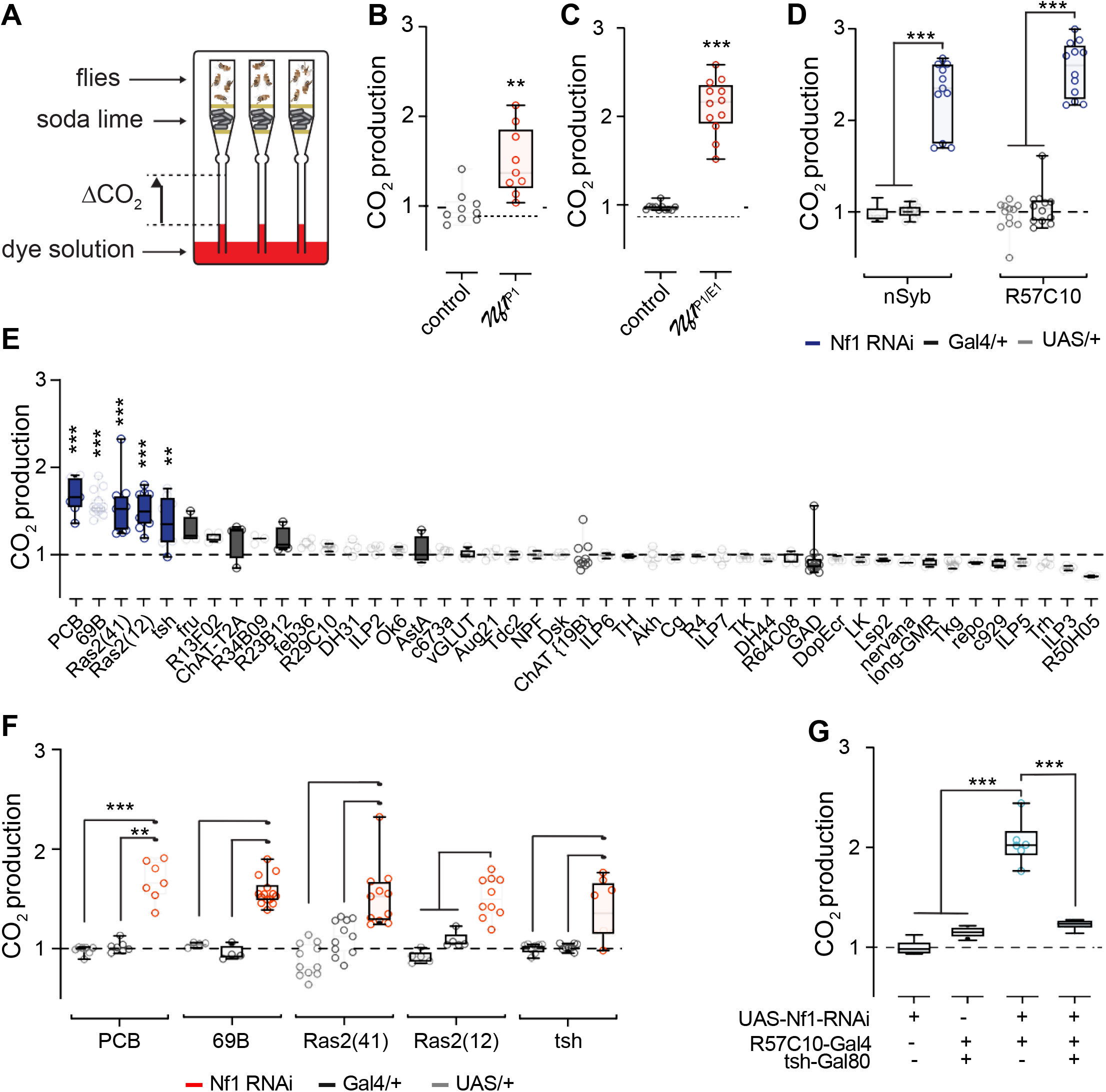
Loss of dNf1 increases CO_2_ production via neuronal mechanisms. (A) Diagram of respirometry apparatus. (B) CO_2_ production in *Nf1*^P1^ mutants and *w*^CS10^ controls, with mutants normalizedto controls. **p < 0.01 (Mann-Whitney, n = 9). (C) Normalized CO_2_ production in heteroallelic *Nf1*^P1/E1^ mutants and matched genetic background controls. ***p < 0.001 (Mann-Whitney, n = 11-12). (D) CO_2_ production when dNf1 was knocked down with RNAi driven by pan-neuronal Gal4 lines: nSyb- and R57C10-Gal4. Gal4/+ and UAS/+ are heterozygous controls. For each driver, data are normalized to the mean of both controls. ***p < 0.001 (Dunn’s test, n = 8-12). (E) Screen for neuronal subsets in which knocking down dNf1 elevates CO_2_ production. Normalized CO_2_ production from each Gal4 line is shown. (F) Significant lines from panel E, showing both RNAi and heterozygous control groups. *p < 0.05, **p < 0.01, ***p < 0.001 (Dunn’s test, n = 7-14). (G) Normalized CO_2_ production when knocking down dNf1 pan-neuronally with R57C10-Gal4, with and without the addition of the tsh-Gal80 repressor. ***p < 0.001 (Sidak, n = 6).

To identify the neurons responsible for the increase in CO_2_ production, we knocked down dNf1 using neuronal subset-selective Gal4 drivers. Drivers were selected that express in neurons that release particular neurotransmitters (e.g., cholinergic neurons [ChAT]), modulate growth and/or metabolism (e.g., insulin-like peptides [ILPs], c673a) (Al-Anzi et al., 2009), or mediate previously characterized dNf1 effects on growth in larvae (e.g., Ras2) (Walker et al., 2006). For each line, the experimental group was compared to the two heterozygous genetic controls(Gal4/+ and UAS/+) within experiments. Among the 45 lines tested, 5 lines exhibited significant elevation of metabolic rate relative to both controls (Fig. 1 E,F and S1E). The positive lines were, in descending order of effect size, PCB, 69B, Ras2(41), Ras2(12), and tsh (Fig. 1F and S1E). The PCB-Gal4, described in more detail below, expresses in a sparse subset of neurons in the nervous system. The 69B and Ras2 drivers express broadly across the nervous system. Within neurons, the tsh-Gal4 line expresses preferentially in the ventral nervous system (VNS), suggesting that neurons in the VNS may be a major contributor to the metabolic phenotype. To further localize the effect, we parsed the brain and VNS with an intersectional approach. We drove the dNf1 RNAi pan-neuronally with R57C10 in one set of flies and suppressed VNS expression with the tsh-Gal80 repressor in another group (Fig. 1G and S1D). The tsh-Gal80 repressor differentially removes expression from neurons in the VNS (Clyne and Miesenbock, 2008). Limiting expression with tsh-Gal80 eliminated the metabolic phenotype, suggesting that dNf1-sensitive neurons are tsh-positive, and may reside in the VNS.

The mapping experiments revealed several major aspects of the dNf1-mediated neuronal control of metabolism. First, the dNf1 effect on metabolic rate was not due to the effects on a selective neurotransmitter system or neuronal subset (notably GABAergic and cholinergic neurons and neurons that release insulin-like peptides). Instead, knocking down dNf1 in relatively broader sets of neurons in the brain and/or VNS was required to drive an increase in metabolic rate. The driver with the most restricted expression pattern that produced a large effect was the PCB-Gal4 driver (Fig. 2A). This driver has a Gal4 enhancer trap in the pyruvate carboxylase (PCB) locus, and it has been previously termed “fatbody-Gal4” due to positive expression in the fat body (Gronke et al., 2003). However, our experiments suggest that the metabolic phenotype is neuronal in origin. Therefore, we tested whether this driver also exhibits neuronal expression. Using the PCB-Gal4 to drive green fluorescent protein (mCD8∷GFP, we fixed and immunostained adult brains and VNS, and found sparse but robust GFP expression in neurons in both regions (Fig. 2A). Given that the driver is not fat body-specific, we propose that it be referred to by its genomic locus (PCB-Gal4) and do so here.

**Figure 2.**
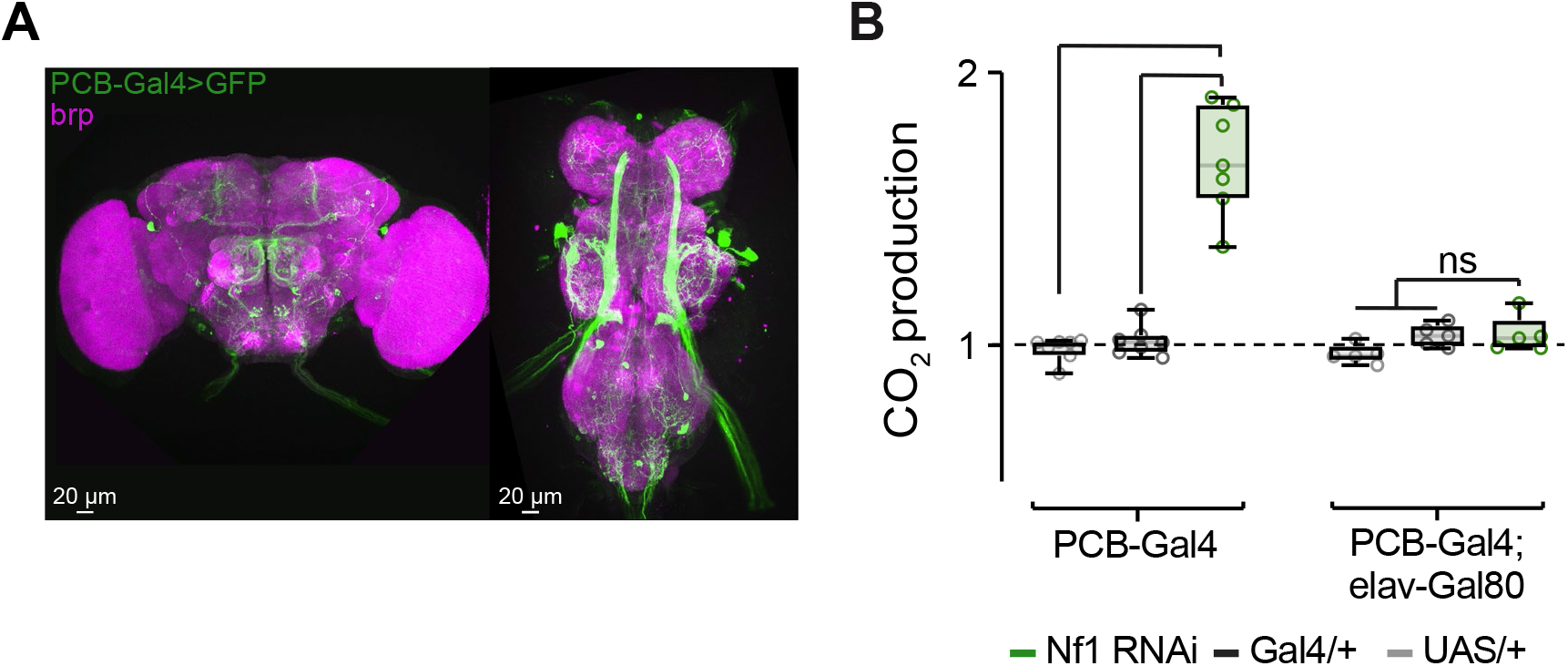
Knocking down dNf1 in a sparse set of neurons increases metabolic rate. (A) Expression of GFP (green) in neurons labeled by the PCB-Gal4 driver. Bruchpilot (brp) was used to counterstain neuropil (magenta). (B) CO_2_ production assayed by respirometry. dNf1 was knocked down in neurons with the PCB-Gal4 driver, with or without the elav-Gal80 repressor. The PCB-Gal4 without Gal80 data set is the same as in Figure 1, shown here for comparison. ns: not significant. **p < 0.01 ***p < 0.001 (Dunn’s test, n = 5-7).

To further examine whether the metabolic effect was due to fat body or neuronal expression, we tested three other fat body-expressing Gal4 drivers: R4, Cg and Lsp2. None of these produced a metabolic phenotype when knocking down dNf1 (Figs. 1E, S2). In addition, we subtracted neuronal expression from the PCB-Gal4 driver with elav-Gal80 (Yang et al., 2009) (Figs. 2B, S3). This eliminated the dNf1-induced metabolic effect, providing additional evidence that neurons in the PCB driver were responsible for the effect. Finally, 69B-Gal4, one of the Gal4 lines with largest effect sizes (Fig. 1 E,F), has previously been shown to be expressed in the central nervous system but not in the fat body (Palgi et al., 2012). Thus, multiple lines of evidence support the interpretation that neuronal expression in the PCB-Gal4 driver, rather than fat body expression, is responsible for the metabolic phenotype. Overall, these data suggest that dNf1 is required in a subset of neurons labeled by the PCB-Gal4 driver for normal metabolic regulation.

To gain further insight into the mechanistic underpinnings of the metabolic phenotype, we turned to stop-flow respirometry, quantifying O_2_ consumed and CO_2_ produced with stop-flow respirometry (Fig. 3A) (Stahl et al., 2017). Mutant *Nf1*^*P1*^ adult flies exhibited increased O_2_ consumption and CO_2_ production across the 24-hr photoperiod relative to controls (Fig. 3 B,D). These differences were significant when the data were binned into both day and night periods (Fig. 3 C,E). Similarly, when dNf1 was knocked down pan-neuronally using nSyb-Gal4, we observed increased O_2_ consumption and CO_2_ production across the circadian photoperiod relative to controls (Fig. 3 G-J). In both cases, the respiratory quotient (RQ), the ratio of CO_2_ eliminated to O_2_ consumed, was significantly reduced in dNf1 loss of function (Fig. 3 F,K). Decreased RQ is consistent with increased utilization of endogenous fat stores (Stahl et al., 2017), suggesting that loss of dNf1 may increase fat utilization, a possibility we consider further below. Overall, these data provide independent support for the role of dNf1 in metabolic regulation, demonstrate that it is consistent across the 24-hr photoperiod, and suggest that it may result from altered fat homeostasis.

**Figure 3.**
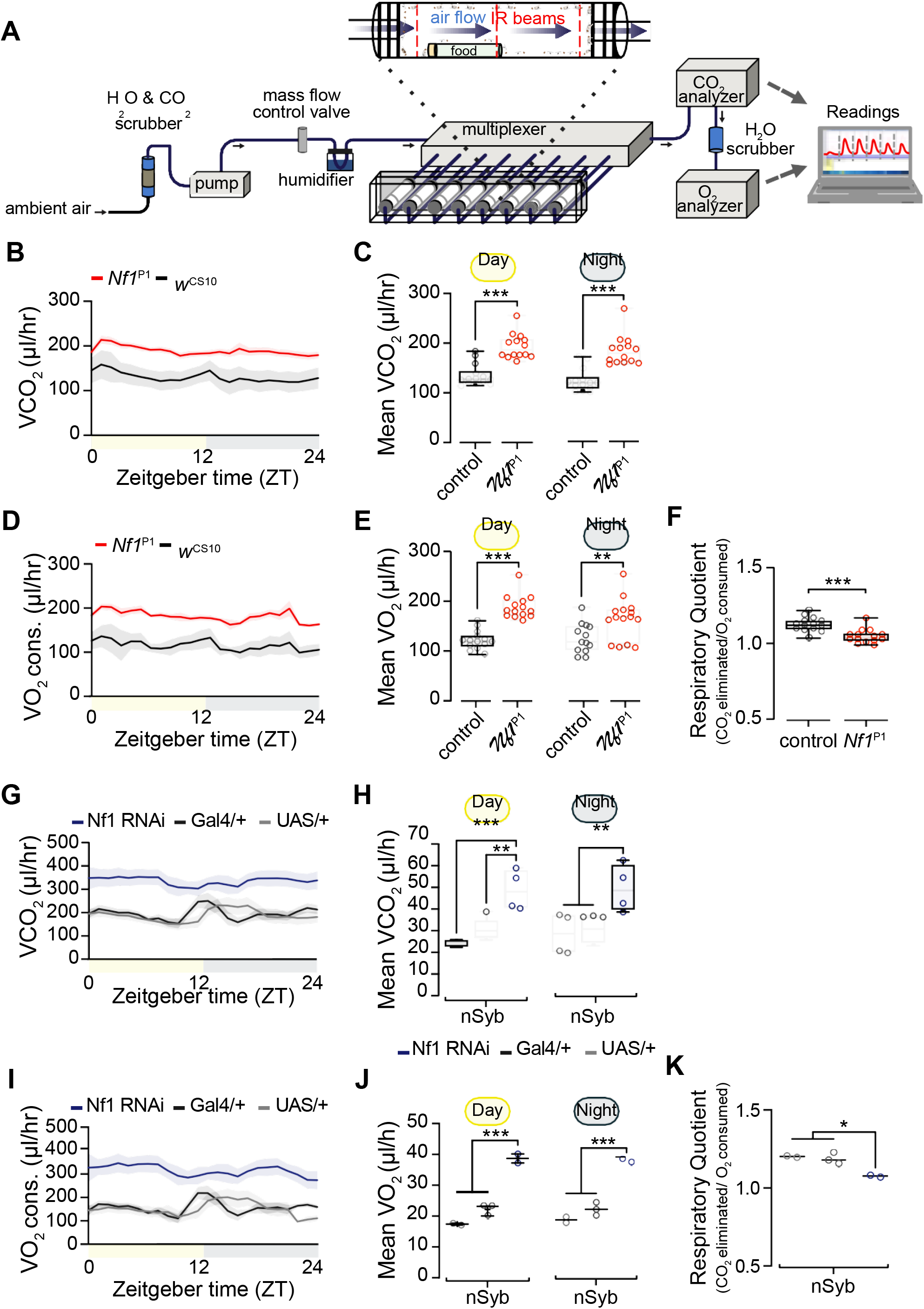
Loss of dNf1 increases metabolic rate across the circadian photoperiod. (A) O_2_ consumption and CO_2_ production were measured with indirect calorimetry. (B) CO_2_ production measured in *Nf1*^P1^ mutants and *w*^CS10^ controls, plotted in 1-hr bins across the circadian photoperiod. (C) Quantification of CO_2_ production during day and night periods. ***p < 0.001 (Sidak, n = 14). (D) O_2_ consumption measured in *Nf1*^P1^ mutants and *w*^CS10^ controls, plotted in 1-hr bins across the circadian photoperiod. (E) Quantification of O_2_ consumption from panel D during day and night periods. **p < 0.01, ***p < 0.001 (Sidak, n = 13-15). (F) Respirometry quotient in *Nf1*^P1^ mutants and *w*^CS10^ controls. ***p < 0.001 (Mann-Whitney; n = 14). (G) CO_2_ production, comparing pan-neuronal nSyb>RNAi with heterozygous controls, plotted in 1-hr bins across the circadian photoperiod. (H) Quantification of CO_2_ production from panel G during day and night periods. **p < 0.01, ***p < 0.001 (Sidak, n = 4-6). (I) O_2_ consumption, comparing pan-neuronal nSyb>RNAi lines with heterozygous controls, plotted in 1-hr bins across the circadian photoperiod. (J) Quantification of O_2_ production from panel I during day and night periods. ***p < 0.001 (Sidak, n = 2-3). (K) Respirometry quotient in nSyb>RNAi lines and heterozygous controls. *p < 0.05 (Sidak, n = 2-3).

### Metabolic regulation is independent of grooming

Loss of dNf1 increases spontaneous grooming (King et al., 2016), which could drive an increase in energy expenditure. To test whether this accounts for the increase in metabolic rate observed here, we knocked down dNf1 and quantified grooming in an open field arena (Fig. 4A). Pan-neuronal knockdown of dNf1 elevated spontaneous grooming (Fig. 4B), as previously reported (King et al., 2016). In addition, we tested the more restricted PCB-Gal4 driver. While knocking down dNf1 with this driver produced one of the largest increases in metabolic rate (Fig. 1 E,F), it did not significantly elevate spontaneous grooming (Fig. 4C). Therefore, dNf1 functions in independent populations of neurons to regulate metabolic rate and grooming. Further, these findings reveal that the elevated metabolic rate observed in *dNf1* mutant flies is not due to changes in grooming.

**Figure 4.**
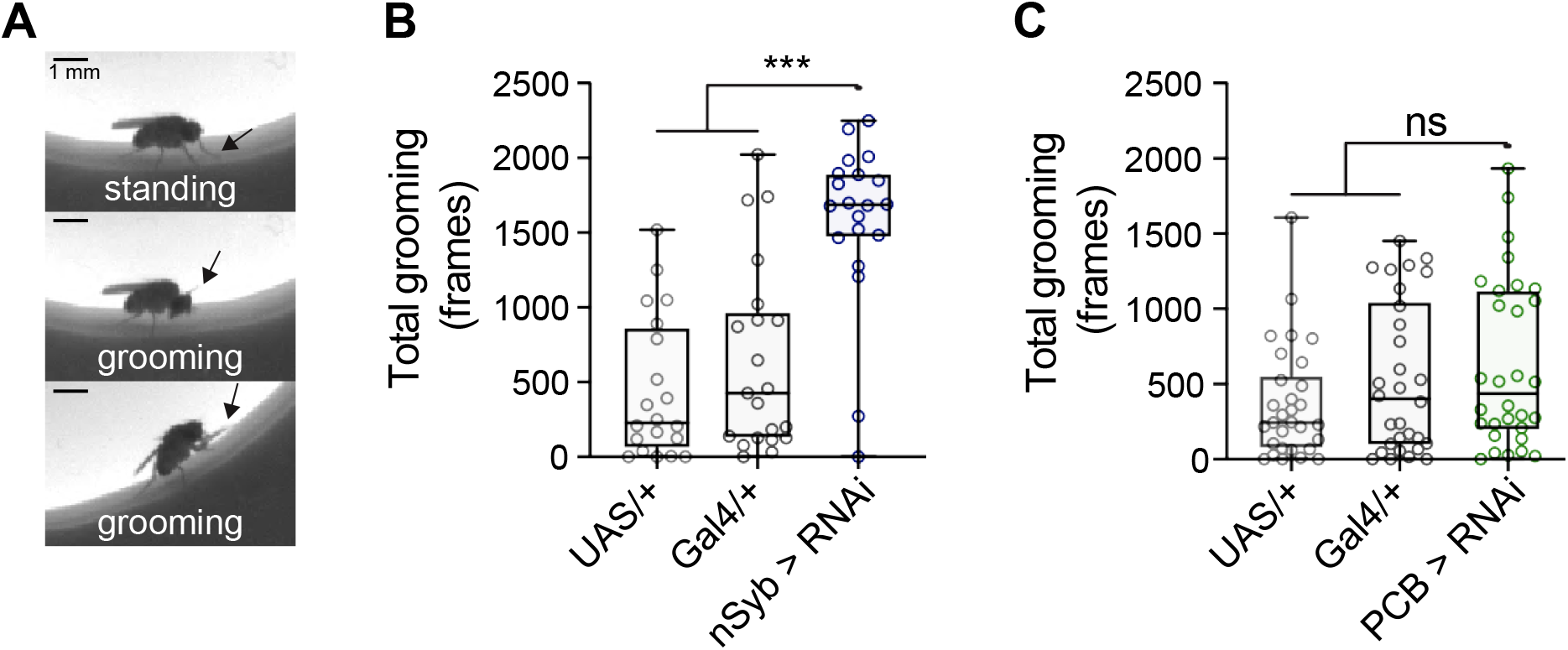
Knockdown of dNf1 in PCB-Gal4+ neurons does not affect spontaneous grooming. Representative images of fly grooming in an open field. Top panel: a fly standing in place (not grooming). Middle panel: a fly grooming its head. Bottom panel: a fly grooming its prothoracic (front) legs. In all panels, an arrow points to the prothoracic leg location. Scale bars = 1mm. (B) Quantification of grooming in flies in which dNf1 has been knocked down pan-neuronally with the nSyb-Gal4 driver. p < 0.001 (Kruskal-Wallis), ***p<0.001 (Dunn; n = 20-21). (C) Quantification of grooming in flies in which dNf1 has been knocked down with the PCB-Gal4 driver. ns: not significant. p = 0.209 (Kruskal-Wallis, n = 30).

### Loss of dNf1 increases starvation susceptibility by decreasing lipid stores via increased lipolysis

Alterations in metabolic rate could be expected to affect lipid storage, and the decreased respirometry quotient suggests that lipid stores may be reduced. To directly test this, we first quantified triglyceride content using coupled colorimetry (Fig. 5A) (Tennessen et al., 2014). Triglyceride stores were significantly reduced in *dNf1* mutants compared to controls (Fig. 5A). Changes in either lipogenesis or lipolysis can decrease triglyceride levels (Fig. 5B). To determine which of these processed is dysregulated, we examined lipid turnover in a pulse-chase experiment (Fig. 5C). Flies were fed radiolabeled ^14^C sucrose for 24 hr. Incorporation of 14C sucrose into fatty acids in the flies was measured with scintillation at 0 hr or 48 hr after returning flies to normal, unlabeled food. Initial incorporation of ^14^C did not differ between controls and *Nf1*^*P1*^ mutants, suggesting that the synthesis of fatty acids (i.e., lipogenesis rate) was unchanged (Fig. 5D). However, after 48 hr, *Nf1*^*P1*^ flies retained significantly less ^14^C than controls, suggesting that the mutants exhibited increased rates of lipolysis. The effect was significant regardless of whether incorporation was normalized to body weight or the number of flies (data not shown). Together, these findings suggest that lipid levels are reduced in *dNf1* mutant flies due to increased lipolysis.

**Figure 5.**
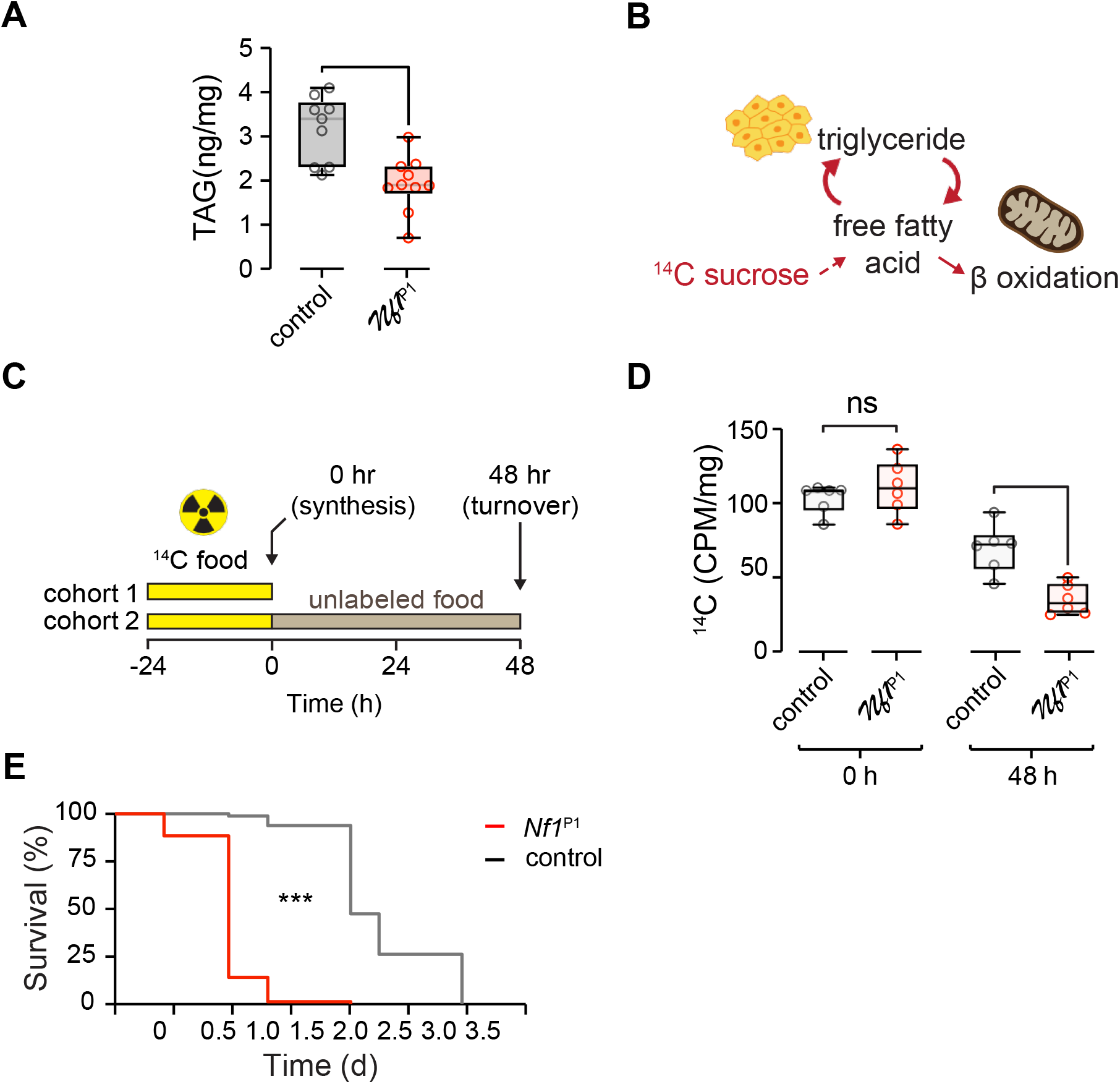
Loss of dNf1 reduces starvation resistance, decreases triglyceride levels, and increases lipid turnover rate. (A) Quantification of total body triglyceride (TAG) content in *Nf1*^P1^ mutants and *w*^CS10^ controls. **p < 0.01 (Mann-Whitney), n = 9-10. (B) Diagram of ^14^C sucrose radiolabeling and lipid turnover. (C) Experimental protocol of ^14^C sucrose radiolabeling protocol for lipid turnover analysis. (D) Quantification of lipid turnover via ^14^C sucrose radiolabeling. Radioactivity was measured as counts per minute (CPM) and normalized to fly weight (mg). ns: not significant, ***p < 0.001 (Sidak; n = 6). (E) Lifespan under starvation, comparing *Nf1*^P1^ mutants (n = 78) and *w*^CS10^ controls (n = 80). *** p < 0.0001 (X^2^(1) = 154.0, Mantel-Cox test).

Triglycerides provide a major source of nutrients in flies that is critical for survival in nutrient-poor conditions. To determine whether the reduced triglyceride levels and increased metabolic rate impact survival in the absence of food, we measured the starvation resistance of *Nf1*^*P1*^ mutants and controls flies. *dNf1* mutants were significantly more susceptible to starvation, succumbing in less than half the time of control flies when housed on a nutrient-free agar media (Fig. 5E). This suggests that loss of dNf1 increases starvation susceptibility, potentially due to the altered lipid homeostasis, which leaves the flies with lower reserves to draw on during caloric restriction.

### Loss of dNf1 increases feeding in adult flies

Feeding and energy stores are homeostatically regulated, and animals often compensate for depleted energy stores by increasing food intake. Since loss of dNf1 increased metabolic rate, decreased lipid stores, and increased lipid turnover via increased lipolysis, energy intake could be increased as a homeostatic compensatory mechanism. To test this, we examined feeding using a capillary feeding assay (Murphy et al., 2017) (Fig. 6A). Food intake was significantly increased in *Nf1*^*P1*^ adult flies relative to controls (Fig. 6B). Knocking down dNf1 pan-neuronally also increased feeding relative to controls (Fig. 6C), confirming the effect and suggesting that it was due to the loss of dNf1 specifically in neurons. We next tested the neuronal circuit requirements for dNf1 in the context of feeding. A selection of Gal4 drivers that labeled different neurochemical subsets, and/or produced metabolic alterations, were used to drive dNf1 RNAi (Fig. 6D). Among these lines, significant elevations in feeding were observed when dNf1 was knocked down pan-neuronally (nSyb), selectively in the VNS (tsh), and in subsets of neurons labeled by the 69B and PCB-Gal4 drivers. No effect was observed when dNf1 was knocked down in GABAergic neurons (GAD) or insulin-like peptides (ILP2, ILP3, ILP5, ILP 6, ILP7) (Fig. 6D). Thus, knocking down dNf1 in the same circuits that elevated metabolic rate (Fig. 1 E,F) also drove an increase in feeding. While it is possible that dNf1 independently regulates both phenotypes via actions in the same circuit(s), we surmise that the increase in feeding is likely a homeostatic response to the increase in metabolic rate and decrease in fat stores caused by loss of dNf1.

**Figure 6.**
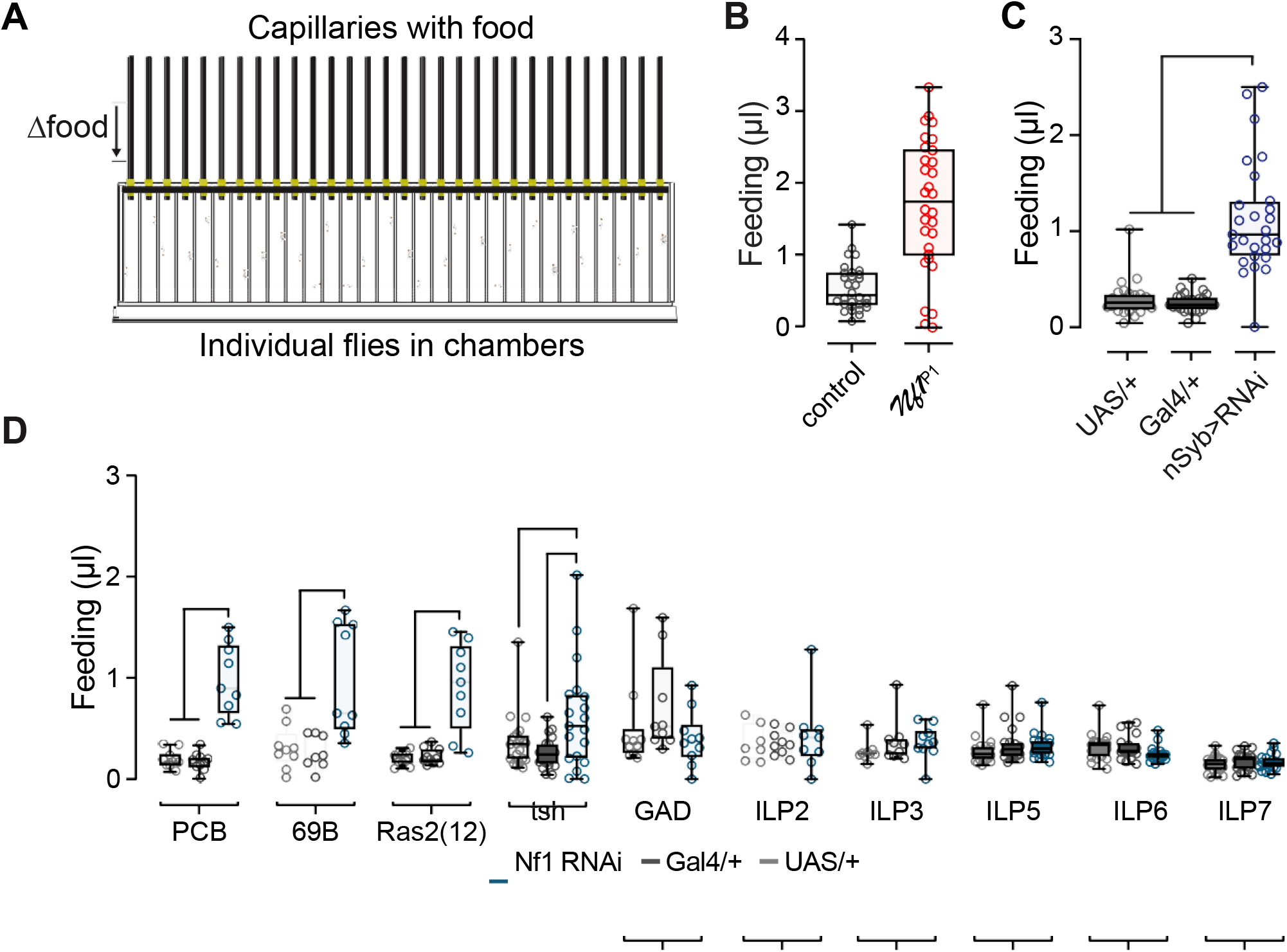
Loss of dNf1 increases feeding. (A) Illustration of the capillary feeding apparatus. (B) Feeding in *Nf1*^P1^ mutants and *w*^CS10^ controls. ***p < 0.001 (Mann-Whitney, n = 30) (C) Feeding with pan-neuronal dNf1 RNAi (driven by nSyb-Gal4) and heterozygous Gal4/+ and UAS/+ controls. ***p < 0.001 (Dunn’s test, n = 24-27) (D) Feeding measured with knockdown of dNf1 in different subsets of neurons with various Gal4 drivers. *p < 0.05, **p < 0.01, ***p < 0.001 (Dunn’s test, n = 7-20).

### Ras GAP-related domain signaling underlies the Nf1 metabolic alterations

While the catalytic RasGAP activity of neurofibromin is located in its central portion, the GRD, other parts of the protein may contain other functions that could mediate cellular effects (Cichowski and Jacks, 2001; Ratner and Miller, 2015). To determine the critical cellular signaling pathway(s) mediating dNf1 effects on metabolic rate, we first tested the role of the GRD. We utilized the UAS/Gal4 system to express dNf1 in flies with the heteroallelic *Nf1^P1/E1^* mutant combination. Pan-neuronal rescue of full-length, wild-type dNf1 restored metabolic rate to control levels (Fig. 7 A,B). In contrast, pan-neuronal expression of full-length dNf1 carrying a missense mutation in the “arginine finger” of the GRD, R1320P, did not rescue the metabolic effect (Fig. 7 A,B). This mutation corresponds to the patient-derived R1276P mutation in *NF1* shown to reduce Ras GAP activity >1000 fold (Klose et al., 1998; Walker et al., 2006). These data suggest that expression of wild-type dNf1 in neurons restores normal metabolic rate and that RasGAP activity is necessary for dNf1 metabolic effects.

**Figure 7.**
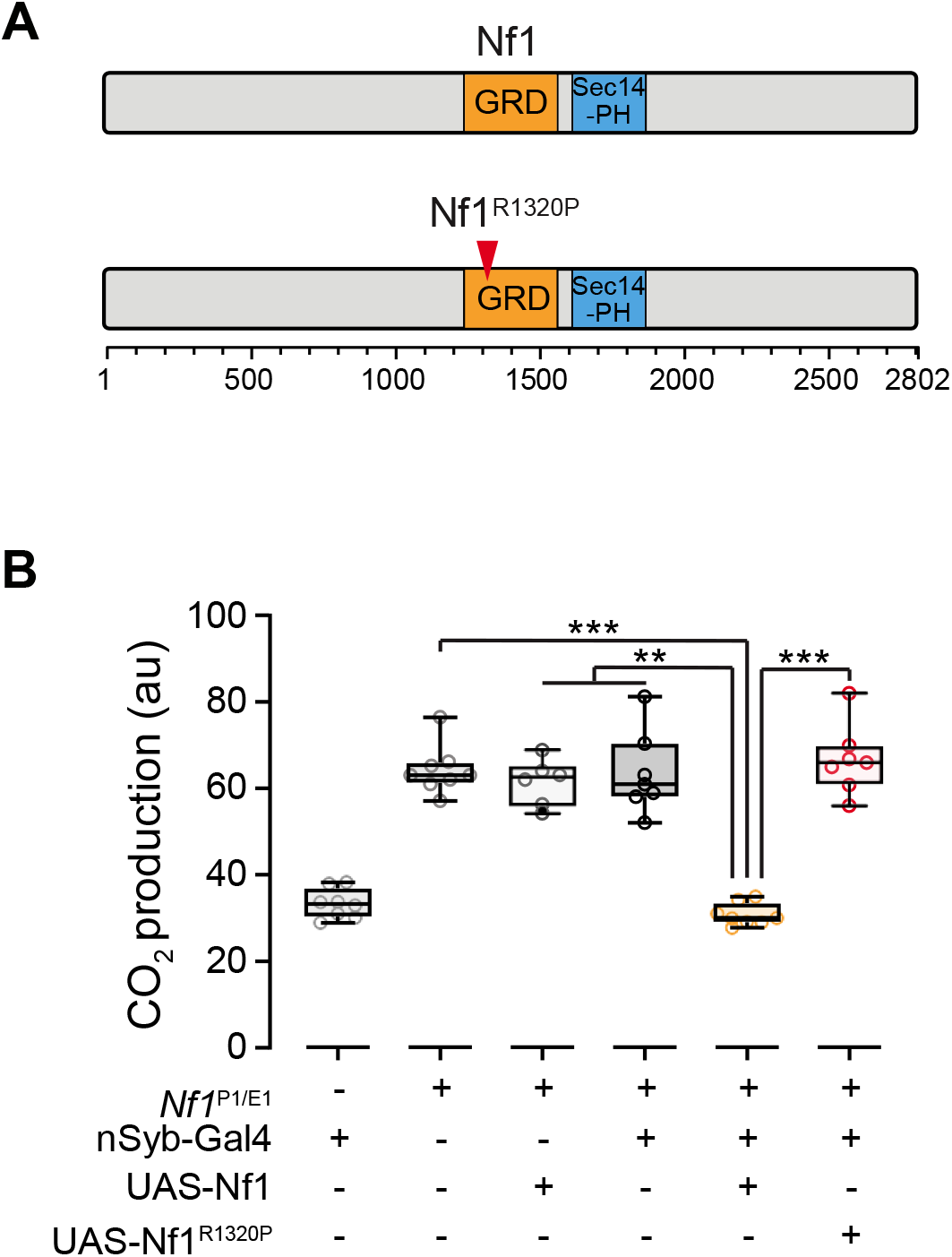
dNf1 metabolic effects are rescued by full-length dNf1 and require a functional GAP-related domain. (A) Major domains in neurofibromin and the site of the R1320P mutation. GRD: GAP-related domain, PH: pleckstrin homology-like domain. (B) Transgenic expression of full-length dNf1 and dNf1^R1320P^ in the heteroallelic *Nf1*^P1/E1^ mutant background. Full length dNf1 rescues CO_2_ production to nSyb-Gal4/+ levels. **p < 0.01 ***p < 0.001 (Dunn’s test, n = 6-8). au: arbitrary units.

## DISCUSSION

This study provides insight into the role of dNf1 in regulating basal metabolic rate via neuronal mechanisms in *Drosophila*. The data support the following major specific conclusions about the role of dNf1 in regulating metabolic rate: 1) loss of dNf1 increases metabolic rate, independent of locomotor changes (grooming), 2) loss of dNf1 function decreases fat stores via increased lipolysis, while increasing food consumption, 3) dNf1 modulates metabolism via a subset of neurons in the VNS labeled by the PCB-Gal4 driver, and 4) dNf1 acts via the GRD to alter metabolic function.

We found that dNf1 plays a central role in regulating metabolic rate via both indirect and stop-flow respirometry. Null genomic mutations and pan-neuronal RNAi knockdown of dNf1 caused significant elevations in metabolic rate. We tested multiple driver lines to determine which cell types are most affected by dNf1 loss of function, finding that dNf1 acts on a relatively narrow set of neurons to modulate metabolic rate. These neurons were not ones commonly implicated in metabolism, feeding, or organismal growth, as knocking down dNf1 in insulin-producing cells, peptidergic neurons, ring gland cells, monoaminergic neurons, etc., produced no effect. It was particularly surprising that loss of dNf1 function in the insulin-producing cells yielded no significant changes in metabolism or feeding, as the insulin-like peptides released by these neurons are functional homologs of insulin and insulin-like growth factors. Furthermore, alterations to such cells affect starvation response and lipid stores (Baker and Thummel, 2007).

A sparse set of neurons in the nervous system, labeled most discretely by the PCB-Gal4 driver, was responsible for the dNf1 effect on metabolism. Due to additional fat body expression with this driver, we verified that the metabolic phenotype was due to loss of neuronal dNf1, using additional Gal4 lines that express in the adult and larval fat body. The fat body serves an important homeostatic function by storing lipids, similarly to white adipose tissue in mammals, and plays a role in the regulation of peripheral tissues under both fed and starved conditions (Baker and Thummel, 2007). Therefore, the lack of metabolic effects with multiple fat body specific drivers was notable. The effects of dNf1 on metabolic rate could theoretically be due to altered energy expenditure within neurons themselves or central control of metabolism. Our experiments implicate the latter, as the PCB-Gal4 driver labels a sparse neuronal population and drives a robust metabolic effect when used to knock down dNf1.

A previous study from our group observed increased grooming activity in *dNf1* mutants and with pan-neuronal RNAi. Increased locomotor activity, including activity such as grooming, might be expected to drive increases in metabolism. However, the dNf1 metabolic and grooming phenotypes were dissociable with the PCB-Gal4 driver, which increased metabolic rate without increasing grooming. Thus, we surmise that the effects of dNf1 on grooming and metabolic rate are caused by distinct neuronal circuits, with relatively little overlap. A thorough analysis of the neuronal circuit driving both phenotypes (particularly grooming) will be necessary to understand how dNf1 independently modulates these behaviors.

The metabolic effects we observed in the *Drosophila* model, combined with the conserved signaling and cellular functionality, suggest that loss of dNf1 could produce metabolic alterations across taxa. Ras is upstream of multiple signaling cascades that influence cellular metabolism, including mTOR, and ERK. mTOR and ERK are key metabolic regulators, exerting effects on gluconeogenesis, protein synthesis, adipocyte differentiation, lipid/cholesterol homeostasis, lipogenesis, and lipolysis (Gehart et al., 2010). Conditional knockout of dNf1 in mouse models, while rapidly lethal, produces metabolic alterations in muscle, including increased triglyceride content and activities of oxidative metabolism enzymes (Sullivan et al., 2014). Further, mouse embryonic fibroblasts lacking dNf1 show changes in basal metabolic rate and mitochondrial bioenergetics (Masgras et al., 2017). These findings, along with data from the present study, suggest the potential for homeostatic regulation of metabolic rate via dNf1-mediated mechanisms.

The most well-characterized biochemical function of neurofibromin is its RasGAP activity contained within the GRD. In the present experiments, transgenic expression of full-length, wild-type neurofibromin restored normal metabolic rate in *dNf1* mutants. However, expression of a dNf1 transgene containing the catalytically dead R1320P mutation failed to rescue. This mutation recapitulates the human patient-derived R1276P mutation that impacts the arginine finger of the GRD, reducing RasGAP activity >1000 fold (Klose et al., 1998). In addition to the GRD, neurofibromin contains other domains with potential functional significance, such as a lipid-binding pleckstrin-homology domain (Fig. 7). Thus, this experiment demonstrated that catalytic activity from the GRD itself was critical for dNf1 modulation of metabolic rate, rather than other putative functions such as lipid binding. Some studies have implicated Ras/GRD function in driving dNf1 phenotypes (Costa et al., 2002; Walker et al., 2006; Williams et al., 2001), while others have identified other signaling roles, for instance on dopaminergic signaling, G protein signal transduction, and cAMP signaling (Brown et al., 2010; Dasgupta et al., 2003; Diggs-Andrews et al., 2013; The et al., 1997; Walker et al., 2013; Wolman et al., 2014; Xie et al., 2016). Our data suggest that the GRD is necessary for dNf1-dependent modulation of metabolic rate, thereby implicating neuronal Ras signaling. The effects of aberrant Ras signaling could include transcriptional regulation, as Ras regulates the activity of multiple transcription factors (Masgras et al., 2017). Further, alterations in these signaling cascades may affect the release of neurotransmitters in ways that produce non-cell-autonomous effects on signaling cascades in other cells (Walker et al., 2006).

Among individuals with NF1, there are subtle alterations in growth, particularly reduced stature (Soucy et al., 2013), as well as reduced body mass index in males (Koga et al., 2016). Alterations in specific metabolites have been reported, though these are sex-specific (Koga et al., 2016). Diabetes mellitus – as well as deaths from noted diabetes complications – is rare in patients with NF1 (Koga et al., 2016). Loss of NF1 increases glycolysis and decreases respiration via mitochondrial ERK signaling, which may play a role in tumor growth (Masgras et al., 2017). Finally, increased resting energy expenditure was recently reported in a cohort of female NF1 patients as well as a decrease in respiratory quotient compared to healthy controls (Souza et al., 2019). Therefore, NF1 has the potential to regulate cellular and organismal metabolism, and alterations in metabolism may contribute to the pathophysiology of NF1. Our data suggest that changes in neuronal metabolic control may be a feature of the cellular and organismal alterations that occur following loss of dNf1.

## ACKNOWLEDGMENTS

We thank Dr. Norbert Perrimon for Tk-gut-Gal4, Dr. Yuh-Nung Jan for ILP7-Gal4 and elav-Gal80, and Dr. Toshihiro Kitamoto for DopEcr-Gal4. Fly stocks obtained from the Bloomington Drosophila Stock Center (BDSC) (NIH P40OD018537) and from the Vienna Drosophila Resource Center (VDRC) were used in this study. Research support was provided by NIH R01 NS097237 (S.M.T), NIH R01 AG045036 (W.W.J.), NIH R01 NS085152 and DC017390 (A.C.K.), and NIH R21 NS096402 and DOD-NFRP W81XWH-16-1-0220 (J.A.W.).

## AUTHOR CONTRIBUTIONS

S.M.T. provided overall project management and funding acquisition. S.M.T. and V.B. designed the project and drafted the manuscript. The manuscript was reviewed and edited by V.B., S.M.T., A.C.K., J.A.W., K.J.C., W.W.J, T.B., B.A.S., E.C.G., and K.R.M. Experimental methodology was performed by V.B., W.W.J., A.C.K., and K.J.C. Experiments were performed by V.B., B.A.S., K.R.M., E.C.G., T.B., S.P., L.B.K., and W.W.J. Project resources were provided by S.M.T., W.W.J., A.C.K., J.A.W., and K.J.C. Experimental and project supervision was provided by S.M.T., W.W.J., A.C.K., K.J.C., and L.B.K.

## METHODS

### *Drosophila* **Husbandry and Stocks**

Flies were cultured on cornmeal/agar food medium according to standard protocol and housed at 25 °C, 60% relative humidity, on a 12:12 light:dark cycle. The *Nf1*^*P1*^ mutation was backcrossed six generations into the *w*^CS10^ genetic background and *w*^CS10^ flies were used as controls. Genetic control for *Nf1*^*P1*^/*Nf1^E1^* trans-heterozygotes is *w*^CS10^/w^iso2;3^. UAS-dNf1-eGFP transgenes were generated using a *dNf1* mini-gene (full-length *dNf1* cDNA corresponding to the RF isoform with addition of introns 9 and 10). The R1320P mutation was created in a wild-type cDNA using the Q5 Site-Directed Mutagenesis Kit (New England Biolabs). Wild-type and R1320P mutant *dNf1* were then subcloned into the pUAST-attB vector with an in-frame C-terminal fusion with eGFP cDNA. Transgenic lines were produced by integrating the constructs at the attp40 site (Rainbow Transgenic Flies Inc.) The dNf1 RNAi line was obtained from the Vienna Drosophila RNAi Center (VDRC #109637), Gal4/+ control crosses consisted of an empty attP control line (VDRC #60100). UAS-dicer2 was included to potentiate the RNAi effect (Dietzl et al., 2007), and was included with the UAS-RNAi (i.e., all experimental and UAS/+ genotypes). Male flies were used for all experiments to prevent egg accumulation in behavioral chambers.

### CO_2_ measurement using respirometry

Respirometers were prepared as previously described (Yatsenko et al., 2014). Briefly, a 1 ml pipette tip and 50μl capillary micropipette were securely glued together. Soda lime was placed into each pipette tip between two foam pieces to avoid contact with flies. 16 respirometers were hung on a custom-made rack in a latch-lid measurement chamber. Flies were anesthetized with CO_2_ and allowed to recover for at least 24 hr before beginning the experiment. Four flies of the same genotype were placed into each pipette using an aspirator and tightly sealed at the top using non-hardening modeling clay. One pipette was left empty in each chamber as a temperature and atmospheric control. The measuring chamber was filled with 500mL red-dye solution made with water-based dye. The latch-lid chamber with flies were left to equilibrate to incubator conditions (25 °C) for 1 hr before starting the experiment. Vacuum grease was applied between the latch-lid and chamber to reduce atmospheric and temperature fluctuations. Images were captured every 15 minutes using a time-lapse image software (PhenoCapture) and analyzed using Fiji, measuring the liquid meniscus position in each capillary. This measurement was subtracted from a reference image and repeated at each time point for 3 hr.

### Indirect calorimetry

Metabolic rate was measured using the Sleep and Activity Metabolic Monitor (SAMM) system at 25 °C through indirect calorimetry using a stop-flow, push-through respirometry system (Sable Systems International). Metabolic rate in group-housed adult flies was measured as previously described (Stahl et al., 2018; Stahl et al., 2017). Briefly, the experimental system assessed baseline CO_2_ levels from an empty chamber to measure CO_2_ production and O_2_ consumption from 25 male adult flies. Air was flushed from each chamber for 50 sec to provide a readout of CO_2_ accumulation and O_2_ consumption over a 10-min period. A H_2_O and CO_2_ scrubber was used to dehumidify and remove CO_2_ from air before being pumped into a mass flow control valve to maintain a consistent flow rate. The air was then re-humidified by passing through a reservoir containing deionized water prior to reaching the behavioral chambers. Each behavioral chamber (70 mm long × 20 mm diameter glass tube) contained 25 adult male flies with a food vial containing 1% agar and 5% sucrose. Flies were allowed to acclimate to system for 24 hr before the start of each experiment. All experimental runs included a food-only control baseline chamber. CO_2_ production was analyzed using a LI-7000 CO_2_/H_2_O Analyzer (LI-COR) and then dehumidified using a H_2_O scrubber before O_2_ consumption was measured Oxilla Dual Absolute Differential Oxygen Analyzer (LI-COR). CO_2_ production and O_2_ consumption recordings were taken every 10 min. Respirometry quotient (RQ) was calculated as volume of CO_2_ eliminated/O_2_ consumed.

### Immunohistochemistry

Five to seven-day old PCB-Gal4 adult nervous systems were dissected in 1% paraformaldehyde in S2 medium and processed as previously described (Jenett et al., 2012). Samples were stained with primary antibodies for 3 hr at room temperature and at 4 °C overnight, followed by secondary antibodies for 3 hr at room temperature and 4 d at 4 °C. Incubations were performed in blocking serum (3% normal goat serum). Samples were mounted in Vectashield (Vector Laboratories) for analysis. The following antibodies were used: rabbit anti-GFP (1:1000, Invitrogen), mouse anti-nc82 (1:50, DSHB), goat anti-rabbit IgG and goat anti-mouse IgG (1:800, Alexa 488 or Alexa 633 respectively, Invitrogen). Leica TCS SP8 confocal microscopes were used to obtain images.

### Total Grooming

Four to six-day old PCB-Gal4 adult males were used to measure total grooming, as previously described (King et al., 2016). Briefly, individual animals were aspirated into an open field area, habituated for 15 min, and a 5-min video recording was recorded for analysis. Videos were manually scored and start and stop frames were recorded for each grooming event. Total grooming was calculated as the sum of all grooming during the 5-min video.

### Starvation survival

Adult male flies (group housed, 20 flies/vial) on 1% (non-nutritional) agar. The percentage of flies alive was measured twice every 24 hrs.

### Coupled colorimetric assay for TAG

Five male adult *w*^CS10^ and *Nf1*^*P1*^ were collected per sample, as previously described (Tennessen et al., 2014). Samples were rinsed in PBS, rapidly homogenized in PBST, and the supernatant was heated at 70 °C for 10 min. The glycerol standard solution (Sigma 2.5 mg/ml triolein equivalent glycerol standard; G7793) was prepared in PBST to generate 0.125, 0.25, and 0.5 mg/ml glycerol standards. Glycerol standards, samples and a PBST blank were then added to two microfuge tubes and the triglyceride reagent (Sigma; T2449) was added to one of the two tubes to free the glycerol backbone. All microfuge tubes were incubated at 37 °C for 60 min. Samples were transferred to a clear, flat bottom 96-well plate and free glycerol reagent (Sigma; F6428) was added to each sample and mixed well. The plate was incubated for 5 min at 37 °C and centrifuged in a swing-bucket rotor. Absorbance was measured at 540 nm. TAG concentration per sample was determined by subtracting the absorbance for the free glycerol in untreated samples from the total glycerol concentration in samples with the triglyceride reagent and then normalized by sample weight. TAG content in each sample is based on the triolein-equivalent standard curve.

### Lipid Analysis

As described (Katewa et al., 2012), lipogenesis and lipolysis were measured in adult flies. Briefly, flies were fed with 5 μCi of ^14^C-sucrose incorporated into standard fly food for 24 hr. After 24 hr, half of the flies were collected and frozen at −80 °C to measure lipogenesis (0 h samples). The remaining flies were transferred to unlabeled standard fly food for 48 hr and then immediately frozen at −80 °C to measure lipolysis (48 h samples). All flies were transferred into pre-weighed 1.7 ml Eppendorf tubes and then weighed to obtain fly mass. Frozen samples were homogenized with 200 μl of 2:1 chloroform: methanol mix for 15 minutes at room temperature and then spun at full speed for 15 minutes at room temperature. The supernatant was recovered and 40 μl of 0.9% (w/v) of NaCl was added. Samples were vortexed and spun gently to separate phases. The upper phase was removed, and the interphase was rinsed twice with 200 μl 1:1 methanol: water solution without mixing. The lower chloroform phase was used for scintillation counting.

### Feeding

Feeding was quantified in adult male flies using a capillary feeding assay, as previously described (Ja et al., 2007). Briefly, individual flies were placed into chambers (46 mm × 7 mm) containing 1% agar at the bottom and a glass capillary at the top. Each glass capillary contained a meniscus labeling dye to track food consumption and an aqueous food solution (2.5% sucrose + 2.5% yeast extract). All flies were habituated to the chamber and liquid diet 72 hr before measuring food intake (glass capillaries were replaced daily). Total feeding was measured over 24 hr and analyzed by Noah.py (Murphy et al., 2017).

### Statistical Analysis

Normality of data was assessed with the D’Agostino-Pearson Test. Hypothesis testing was carried out with Student’s t (parametric), one/two-way ANOVA with post-hoc Sidak (parametric), Mann-Whitney (nonparametric), Kruskal-Wallis with post-hoc Dunn (nonparametric), or Kaplan-Meier with Mantel-Cox (Log-rank) tests. All statistical analyses were performed using GraphPad Prism 8.1.2.

