## Supplemental Files for "Neurofibromin regulates metabolic rate via neuronal mechanisms in *Drosophila*"

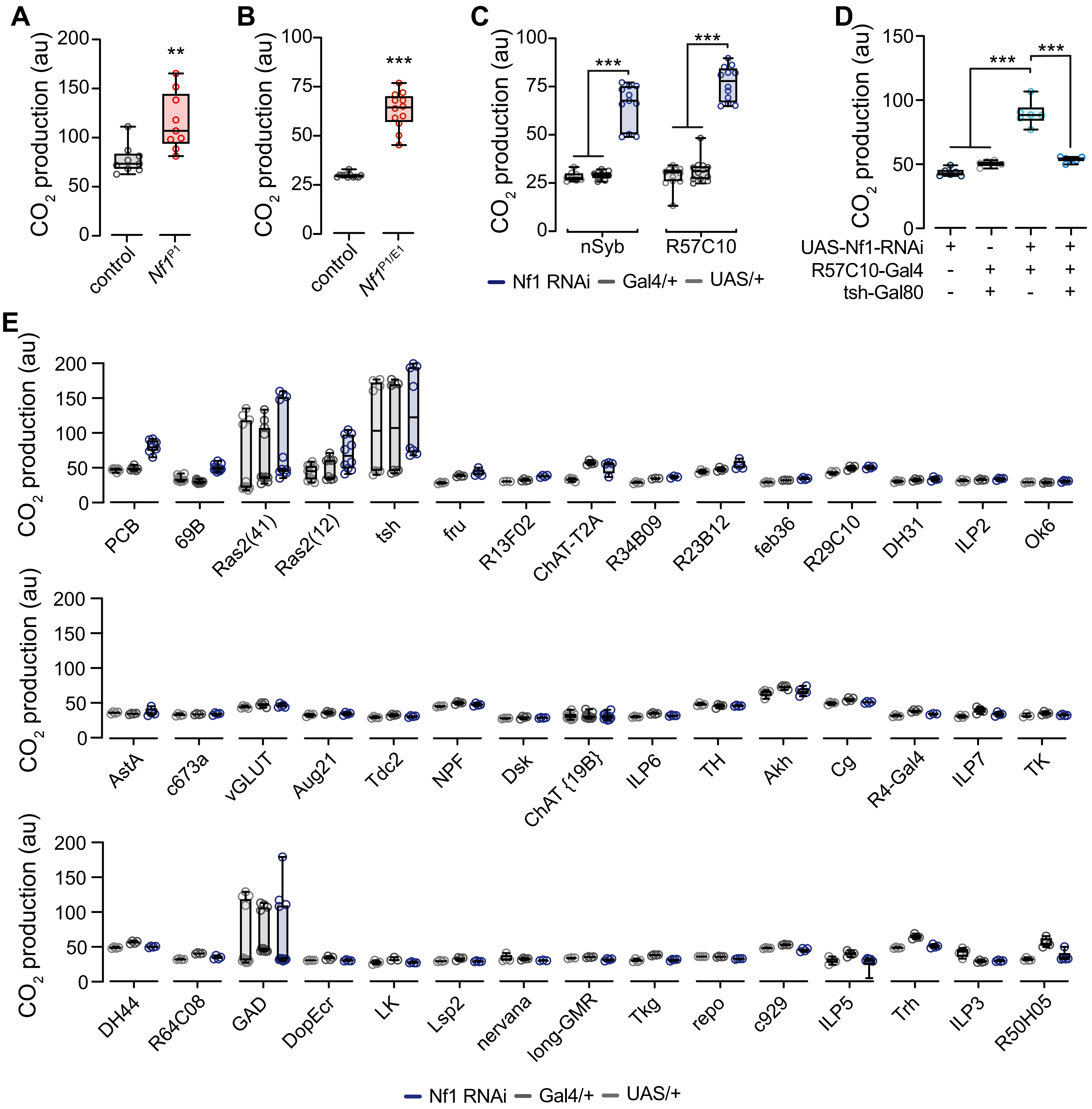


**Figure S1.** Related to Figure 1. (A) $CO_2$ production in *Nf1*P1 mutants and *w*CS10 controls. **p < 0.01 (Mann-Whitney, n = 9). (B) $CO_2$ production in *N1*P1/E1 heteroallelic mutants compared to matched genetic background controls. ***p<0.001 (Mann-Whitney, n = 11-12). (C) $CO_2$ production when dNf1 was pan-neuroanlly knocked down with RNAi using Gal4 lines: nSyb- and R57C10-Gal4. Gal4/+ and UAS/+ are heterozygous controls. ***p<0.001 (Dunn's post-hoc, n = 8-12). (D) Pan-neuronal dNf1 knockdown with R57C10-Gal4, with and without the tsh-Gal80 repressor. ***p<0.001 (Sidak, n = 6). (E) Raw data from the screen for neuronal subsets in which Nf1 knock down elevates $CO_2$ production. au: arbitrary units.

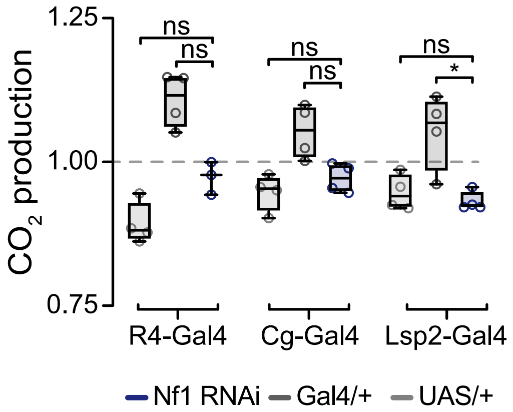


**Figure S2.** Related to Figure 1. $CO_2$ production, assayed by respirometry. dNf1 knockdown with RNAi driven by fat body specific Gal4 lines: R4-, Cg-, and Lsp2-Gal4. Gal4/+ and UAS/+ are heterozygous controls. For each driver, data are normalized to the mean of both controls. ns: not significant, *p < 0.05 ([R4-Gal4, Nf1 RNAi vs. Gal4/+: p = 0.39; Nf1 RNAi vs. UAS/+: p = 0.74], [Cg-Gal4, Nf1 RNAi vs. Gal4/+: p = 0.07; Nf1 RNAi vs. UAS/+: p >0.99] [Lsp2-Gal4, Nf1 RNAi vs. Gal4/+: *p = 0.02; Nf1 RNAi vs. UAS/+: p >0.99]) (Dunn’s, n = 3-4 per genotype).

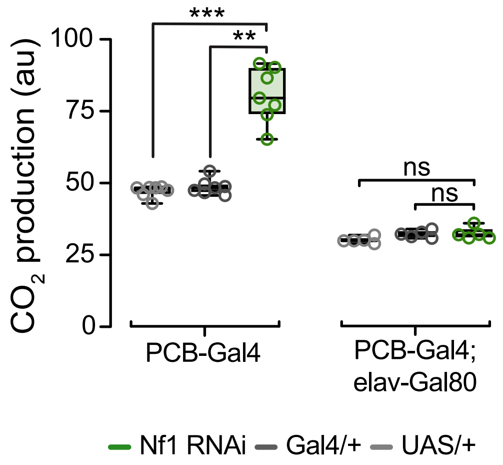


**Figure S3.** Related to Figure 2. $CO_2$ production, assayed by respirometry. dNf1 knockdown in neurons with PCB-Gal4, with or without the elav-Gal80 repressor. ns: not significant, $^{**}p < 0.01$, $^{***}p < 0.001$. (Dunn's, n= 5-7). au: arbitrary units.
